## Supplementary material for "Accurate detection of shared genetic architecture from GWAS summary statistics in the small-sample context": Text S1

Thomas W. Willis    Chris Wallace

**Table S1. Summary statistics for timing benchmarks for candidate bivariate ecdf algorithms used in the GPS permutation procedure.** All values are given in seconds (s).

| ecdf algorithm | Minimum (s) | 1st quartile (s) | Median (s) | Mean (s) | 3rd quartile (s) | Maximum (s) |
|---|---|---|---|---|---|---|
| naive | 4317.7 | 4570.3 | 4759.2 | 4710.8 | 4793.6 | 4956.6 |
| Langrené and Warin | 8.8 | 10.0 | 10.5 | 10.4 | 10.9 | 11.7 |
| Perisic and Posse | 3.8 | 4.9 | 5.0 | 5.0 | 5.2 | 5.6 |

**Table S2. The number of cases and controls, and case-control ratio in the five sample configurations used in our simulation studies.**

| No. of cases | No. of controls | Case-control ratio |
|---|---|---|
| 500 | 10,000 | 0.05 |
| 1,000 | 10,000 | 0.1 |
| 5,000 | 10,000 | 0.5 |
| 10,000 | 10,000 | 1 |
| 100,000 | 100,000 | 1 |

**Table S3. The number of SNPs included in each analysis.** GPS-Exp, GPS-GEV, and Hoeffding's test were applied to LD-pruned sets which varied with respect to the value of the $r^2$ parameter used in pruning and which is given in the header. For LDSC we report the number of SNPs used in regression after merging the real or simulated data set with the panel of precomputed LD scores (Text S2). For SumHer we report the number of SNPs after merging the real or simulated data set with taggings generated from Phase 3 1kGP data; this number is reported in SumHer log files as the number of predictors for which summary statistics were available (Text S2). We did not analyse the single-chromosome simulated data with LDSC or SumHer nor did we perform whole-genome analyses with $r^2 = 0.5$ or 0.8.

| Data set | LDSC | SumHer | $r^2 = 0.2$ | $r^2 = 0.5$ | $r^2 = 0.8$ |
|---|---|---|---|---|---|
| UKBB with MHC | 1,178,572 | 9,487,152 | 591,015 | 1,524,514 | 2,722,772 |
| UKBB without MHC | 1,171,710 | 9,352,179 | 586,552 | 1,511,656 | 2,696,810 |
| simGWAS whole-genome simulations | 1,118,361 | 8,015,936 | 525,150 | - | - |
| simGWAS chromosome 1 simulations | - | - | 42,537 | 97,256 | 165,910 |

**Table S4. Details of the two single-chromosome simulation regimes.**

| Name | Odds ratio(s) | No. of causal variants | No. of shared causal variants |
|---|---|---|---|
| Large-effect (chromosome 1) | 1.2 | 20 | 5, 10, 15, 20 |
| Small-effect (chromosome 1) | 1.05 | 60 | 15, 30, 45, 60 |

**Table S5. Details of the exemplary GWAS data sets.** 'R5' denotes 'Data Freeze 5', a particular iteration of publicly released FinnGen summary statistics.

| Trait | Category | No. of cases | No. of controls | Collection | Citation |
|---|---|---|---|---|---|
| lupus | immune | 5,201 | 9,066 | - | [2] |
| type 1 diabetes | immune | 18,942 | 501,368 | - | [4] |
| Crohn's disease | immune | 12,194 | 28,072 | - | [5] |
| ulcerative colitis | immune | 12,366 | 33,609 | - | [5] |
| rheumatoid arthritis | immune | 19,234 | 61,565 | - | [6] |
| eczema/dermatitis | immune | 20,052 | 198,740 | FinnGen (R5) | [1] |
| hypothyroidism | immune | 26,064 | 192,728 | FinnGen (R5) | [1] |
| hayfever | immune | 27,419 | 416,137 | Pan-UKBB | [3] |
| asthma | immune | 56,065 | 389,965 | Pan-UKBB | [3] |
| cardiomyopathy | non-immune | 11,400 | 175,752 | FinnGen (R5) | [1] |
| endometriosis | non-immune | 8,288 | 68,969 | FinnGen (R5) | [1] |
| macular degeneration | non-immune | 3,794 | 419,170 | Pan-UKBB | [3] |
| glaucoma | non-immune | 8,591 | 210,201 | FinnGen (R5) | [1] |
| leiomyoma | non-immune | 18,060 | 105,519 | FinnGen (R5) | [1] |
| irritable bowel syndrome | non-immune | 11,159 | 425,748 | Pan-UKBB | [3] |
| cholelithiasis | non-immune | 19,350 | 417,176 | Pan-UKBB | [3] |
| osteoarthritis | non-immune | 39,442 | 404,702 | Pan-UKBB | [3] |
| hypercholesterolaemia | non-immune | 47,737 | 394,138 | Pan-UKBB | [3] |

**Table S6. Median genetic correlation estimates by group and sample size as computed from the exemplary data sets.** 'Mixed' pairs were those with one immune and one non-immune disease or two non-immune diseases. $\hat{r}_g$ denotes estimated genetic correlation. 'No. of cases' gives the smaller number of disease cases in each pair of case-control GWAS. We categorised pairs by their smaller case number in the UKBB collection, not the exemplary collection.

| Group | No. of cases | No. of pairs | Median $\hat{r}_g$ |
|---|---|---|---|
| Immune/immune | $\leq$ 2,000 | 21 | 0.07 |
| Immune/immune | $>$ 2,000 | 15 | 0.15 |
| Immune/immune | Any | 36 | 0.09 |
| Mixed | $\leq$ 2,000 | 54 | 0.11 |
| Mixed | $>$ 2,000 | 63 | 0.07 |
| Mixed | Any | 117 | 0.09 |
