## Supplementary material for "Accurate detection of shared genetic architecture from GWAS summary statistics in the small-sample context": Text S2

### Accurate detection of shared genetic architecture from GWAS summary statistics in the small-sample context
### Text S2: Materials and methods

Thomas W. Willis Chris Wallace

#### LD reference panel

We estimated the SNP-SNP genotype squared correlation coefficient $r^2$ using haplotype data for 503 European individuals from the Phase 3 release of the 1kGP. Using the parental identifiers supplied in the `ped` sample metadata file, we removed five individuals with one or both parents present in the cohort (sample identifiers `HG00155`, `NA07048`, `NA10847`, `NA10851`, and `NA12878`) to ensure all members of the cohort were unrelated. We subjected the data to quality control using `PLINK v2.00a2.3LM`[1,2], retaining only those SNPs with a MAF $\geq 0.005$, a Hardy-Weinberg equilibrium test p-value > 1e-50, and a missingness rate $\leq 0.1$.

For both UKBB and simulated data sets we carried out LD pruning with `PLINK` to produce filtered data sets for use with Hoeffding's test and both GPS tests.

We applied Li et al's choice of $r^2 = 0.2$ in our whole-genome simulations before examining test performance with $r^2 = 0.2$, 0.5, and 0.8 in single-chromosome simulations (Verification and comparison). We found the GPS tests were most powerful for $r^2 = 0.8$ and proceeded with this value for our analysis of UKBB data sets.

The LD-pruned SNP set for use in whole-genome simulated GWAS data sets was generated by first filtering the 1kGP SNPs to retain those with MAF $\leq 0.01$ in the European cohort (see 'Simulated GWAS data sets'), then pruning this set with $r^2 = 0.2$. The UKBB LD-pruned sets were generated by first merging the 1kGP SNPs with the set of SNPs present in the UKBB data sets, then pruning the intersection of these two sets with $r^2 = 0.8$. For all pruning procedures, we specified a 1000kbp window and a step size of 50bp. We tabulate the number of SNPs remaining after the LD pruning of each data set in Table S3.

#### UK Biobank GWAS data sets

We downloaded UKBB GWAS summary statistics from the Neale lab's data repository, in particular the 'GWAS round 2' results released in August 2018 (`www.nealelab.is/uk-biobank`).

These data sets each contained 13,791,467 variants. For analyses in which the MHC was removed from these data sets, we removed all variants in the base pair interval 24,000,000-45,000,000 on chromosome 6, leaving 13,632,572.

The Neale lab included a total of 361,194 biobank samples in their GWAS analyses but case/control status with respect to each (binary) phenotype was not defined for all samples (`http://www.nealelab.is/uk-biobank/faq`), such that the sum of cases and controls varies across our chosen phenotypes (Table 2).

When estimating $\rho$, the correlation induced between marginal SNP effect estimates for a pair of traits by sample overlap in the GWAS of each trait, we were unable to determine the exact sample overlap in GWAS from the Neale collection as individual sample identifiers were not released. We instead used a more recent release of the UKBB data, downloaded in April 2023, to count the overlap of cases and controls between traits and thence calculated $\rho$.

#### Exemplary GWAS data sets

We surveyed publicly available GWAS of our 18 phenotypes and selected those with the largest number of cases in order to provide the most precise genetic correlation estimates possible (Table S5). FinnGen data sets were obtained from Data Freeze 5 ('R5'). Pan-UKBB[6] data sets were obtained as directed on the project website (`https://pan.ukbb.broadinstitute.org/downloads`). We obtained summary statistics for the GWAS of lupus, type 1 diabetes, ulcerative colitis, and rheumatoid arthritis from the NHGRI-EBI GWAS Catalog[7].

The majority of exemplary data sets came from the Pan-UKBB and FinnGen collections, a fact which raises the issue of sample overlap discussed above and in the main body of the text. Both the Pan-UKBB and FinnGen collections share controls and, to a more limited extent, cases between their GWAS. In addition, there is substantial overlap in cases and controls between the GWAS of the 2018 UKBB collection (used as our main 'test set') and the GWAS of the Pan-UKBB collection. Whilst we conduct no joint analyses of data sets from these two collections, insofar as the Pan-UKBB GWAS are not truly 'out-of-sample' with respect to the 2018 UKBB GWAS, they are not ideal as a point of comparison for the latter as the Pan-UKBB correlation estimates will not be independent of the GPS-GEV, GPS-Exp, Hoeffding's, SumHer, and LDSC test statistics obtained from the 2018 UKBB GWAS. We balanced these shortcomings of our exemplary data sets against the advantage of obtaining the largest possible GWAS to provide the most precise estimates of the genetic correlation. The UKBB/Pan-UKBB sample overlap is somewhat mitigated by our using Pan-UKBB GWAS for only 7 of 18 traits and (as noted above for the UKBB collection), the impact of the resulting between-trait effect estimate correlation is expected to be small.

We used version 5.3.2 of the `GWAS_tools` pipeline (`https://github.com/GRealesM/GWAS_tools`) to process each study in order to obtain all summary statistics in a uniform format. We computed genetic correlation estimates between phenotypes using SumHer in the manner described below. For Pan-UKBB data sets, we used summary statistics from analyses which included only European individuals.

The hypertension and high cholesterol GWAS referenced in Text S2 as examples of large biobank-derived GWAS were identified from the Pan-UKBB phenotype manifest made available on the project website (`https://pan.ukbb.broadinstitute.org/downloads`). Hypertension and high cholesterol had `coding` column values of 1065 and 1473, respectively. In identifying these as the largest and second-largest case-control GWAS, we searched rows with a 'categorical' `trait_type` value and excluded non-disease and medical intervention phenotypes as well as excessively broad aggregate phenotypes which contained disparate diseases.

#### The GPS tests

The GPS-Exp p-value was obtained for the test statistic $D$ as the lower tail probability of the value $D^{-2}$, assumed to follow a standard exponential distribution as per Equation 2. We computed this p-value using `pexp` from the `R` `stats` package.

We fitted a GEV distribution to the null realisations of the GPS generated with our permutation procedure using the functions `fgev` and `fitdist` from the `R` packages `evd`[8] and `fitdistrplus`[9], respectively. The p-value was obtained from the fitted GEV cdf as the upper tail probability of the test statistic $D$.

With respect to the bivariate ecdf algorithms discussed in the main text, we used the implementation of Langrené and Warin's algorithm from the STochastic OPTimization library [sic][10] and developed our own implementation of the bivariate ecdf algorithm of Perisic and Posse.

To study the dependence of the fitted GEVD's location, shape, and scale parameters on the number of SNPs in the LD-pruned set, we randomly downsampled the LD-pruned UKBB SNP set to obtain smaller subsets on which to evaluate the GPS statistic.

#### Hoeffding's test

Hoeffding's test evaluates a null hypothesis $H_0$ of the independence of two random variables, $X$ and $Y$; this hypothesis can be expressed in terms of their joint and marginal cdfs as $H_0 : F_{X,Y} = F_X F_Y$. The test is based upon the following measure $D$ of the deviation of the bivariate cdf $F_{X,Y}$ from its assumed form under the null:

$$D = \int (F_{X,Y}(x, y) - F_X(x)F_Y(y))^2 dF(x, y). \tag{1}$$

We computed the test statistic for GWAS p-values using the `hoeffding.D.test` function from the `R` package `independence`[11].

#### Simulated GWAS data sets

We simulated GWAS summary statistics using the `R` package `simGWAS v0.2.0-4`[12]. Following the package vignette, we used haplotypes from European individuals from the Phase 3 1kGP data referenced above and included only those variants with MAF $\geq 0.01$. This yielded 8,995,296 variants in a simulated whole-genome data set and 525,150 after LD pruning with a window size of 1000kb, a step size of 50, and an $r^2$ threshold of 0.2 as described above (Table S3). An exception to this approach was made with the single-chromosome simulations in which we explored the performance of the tests with $r^2 = 0.2$, 0.5, and 0.8. We partitioned the genome into 1,625 approximately independent LD blocks using `ldetect`[13] and simulated summary statistics in a blockwise manner in order to reduce the computational burden of simulation. We simulated at most one causal variant per LD block and selected, arbitrarily, the 2000th SNP in each LD block as the causal variant. The tiny-effect regime was an exception to this approach: to accommodate 1,000 causal variants located in disjoint LD blocks between pairs of simulated GWAS in the case with a zero-valued genetic correlation, we first randomly chose the LD blocks to contain causal variants in each GWAS then randomly selected two SNPs in each of these blocks as causal variants. We ran simulations under six regimes (Table 1). We also varied the number of cases and controls across five sample configurations (Table S2). As cases are typically in far shorter supply than controls in case-control GWAS, we sought to emulate this by choosing a greater number of controls, 10,000, than cases for the first three sample configurations where the number of cases was less than 10,000. For the next two configurations with 10,000 and 100,000 cases, respectively, we chose a matching number of controls.

For the simulations in which we explored the effect of different values of $r^2$ on test performance, we simulated only chromosome 1 rather than the whole genome (Table S4).

To induce a particular genetic correlation, we varied the number of causal variants shared between a pair of simulated GWAS. The number of causal variants per GWAS was kept constant in order to keep heritability per simulated trait constant. We simulated 400 replicate GWAS data sets for each sample configuration and genetic correlation, giving 200 replicate pairs of GWAS for each case number-correlation pair. For a given replicate pair, case number, and specified correlation, the identity of the LD blocks which were to contain causal variants was randomly chosen.

#### Simulated data sets with sample overlap

To estimate the effect of sample overlap on type 1 error rate in the UKBB analyses, we simulated a pair of test statistics from 40,800 independent SNPs (the approximate number of SNPs after LD pruning chromosome 1 with a threshold of $r^2 = 0.2$) where test statistics $Z_1, Z_2$ came from a mixture distribution:

$$\begin{bmatrix} Z_1 \\ Z_2 \end{bmatrix} \sim N\left( \begin{bmatrix} \mu_1 \\ \mu_2 \end{bmatrix}, \begin{bmatrix} 1 & \rho \\ \rho & 1 \end{bmatrix} \right)$$

where

$$\mu_i \begin{cases} = 0 & \text{SNP i not associated} \\ \sim N(\mu_z, \sigma_z^2) & \text{SNP i associated} \end{cases}$$

for a range of $\mu_z$ and $\sigma_z^2$ to simulate lowly- or highly-powered studies. We assumed 40,000 SNPs were not associated, 400 were associated in study 1 but not study 2, and 400 were associated in study 2 but not study 1.

#### LD score regression

We estimated the genetic correlation using the LDSC software as directed in the LDSC GitHub repository's wiki (`https://github.com/bulik/ldsc/wiki`). When estimating the heritability from simulated summary statistics, we constrained the intercept to one as the simGWAS simulation model does not admit confounding. The intercept for genetic covariance estimation was allowed to vary. We used precomputed LD scores for SNPs common in European populations (MAF $\geq 0.05$) from the LDSC repository wiki (`https://data.broadinstitute.org/alkesgroup/LDSCORE/eur_w_ld_chr.tar.bz2`). Use of only common SNPS is recommended by LDSC's authors for routine applications in order to avoid the upward bias in heritability estimates that can be introduced by the extrapolation of per-SNP heritability among common variants to rarer ones (`https://github.com/bulik/ldsc/wiki/LD-Score-Estimation-Tutorial`). Use of this MAF-restricted LD score panel accounted for the far smaller number of SNPs we used in LDSC analyses (Table S3). In spite of this, LDSC's performance was largely comparable to SumHer's, its closest comparator in terms of method and purpose, across simulated and real data sets.

#### SumHer

We downloaded the prebuilt LDAK binaries, version 5.2, from the package website (`https://dougspeed.com/ldak/`). We generated SNP taggings using Phase 3 1kGP data from European individuals and the 'LDAK-Thin' model as described on the LDAK website (`https://dougspeed.com/calculate-taggings/`). We estimated genetic correlation with the `SumHer` component of the LDAK package and the default 'SumHer-GC' model, which assumes that confounding inflation is multiplicative. The SumHer-GC model assumes the LDAK heritability model, not the LDAK-Thin model; the authors nonetheless recommend use of the latter owing to its simplicity and their observation that genetic correlation estimates are generally insensitive to the choice of heritability model (`http://dougspeed.com/genetic-correlations/`). We excluded SNPs which explained more than 1% of the phenotypic variance as per the authors' recommendation as analyses can be sensitive to large-effect loci. `SumHer` does not provide for hypothesis testing of the genetic correlation. We performed a chi-squared test against a null hypothesis of no genetic correlation (i.e. $H_0 : r_g = 0$) by dividing the genetic correlation estimate by its standard error, then squaring this quantity to obtain a test statistic. We used the chi-squared survival function `chi2.sf` from the `scipy.stats` Python package to perform the test.

#### Timing benchmarks

We benchmarked three candidate bivariate ecdf algorithms for use in computing the GPS statistic. This choice was important as estimation of the bivariate ecdf is by far the most demanding step in this computation. The naive algorithm uses two nested loops to determine for each p-value pair $(u, v)$ the size of the set $\{(u_i, v_i)|u_i < u, v_i < v\}$ and has a time complexity of $\mathrm{O}(n^2)$ in the number of p-value pairs $n$ in the input. Both Langrené and Warin's divide-and-conquer algorithm and the order statistic tree-based algorithm of Perisic and Posse have a time complexity of $\mathrm{O}(n \log n)$.

To generate timing data, we recorded the wall clock time for the computation of the GPS statistic for a single data set, replicating this 100 times for each algorithm. The input data set comprised a set of LD-pruned, simulated whole-genome summary statistics containing 525,150 SNPs. LD pruning was conducted with a window size of 1000kb, a step size of 50, and an $r^2$ threshold of 0.2. The `computeGpsCLI` program was used to record running time by specifying the `timingFile` command-line argument and is available in the `gps_cpp` repository. The program was invoked so as to use a single thread in each instance.
