## Supplementary material for "Accurate detection of shared genetic architecture from GWAS summary statistics in the small-sample context": Text S3

### Accurate detection of shared genetic architecture from GWAS summary statistics in the small-sample context
### Text S3: Type 1 error and sample size for the GPS tests

Thomas W. Willis Chris Wallace

We observed that the type 1 error rate of the GPS-Exp and GPS-GEV tests fell as the number of cases increased from 5,000 to 100,000. Fig S11 depicts the observed type 1 error rates with the inclusion of two additional sample sizes, 50,000 and 250,000 cases and controls. We omitted these sample sizes from the main body of the text for the sake of brevity and the unrealistic magnitude of the second for a case-control GWAS, but their inclusion helps to confirm the trend of a falling type 1 error rate for the GPS-Exp and GPS-GEV tests.

We examined the role of the GPS-GEV permutation procedure in this phenomenon. Whereas the permutation procedure was used to generate many permuted null realisations of the GPS statistic from a single given data set, we also obtained null realisations of the GPS statistic through direct simulation as part of our comparative whole-genome simulation study: the generation of 200 replicate simulations with a genetic correlation of 0 for every sample size provided us with 200 'empirical' null realisations of the GPS statistic for each sample size and simulation regime.

As the number of cases and controls increased, the number of large values of the GPS statistic decreased for empirical, but not permuted, data sets (Fig S12). The permutation procedure failed to recapitulate the behaviour of empirical data at higher sample sizes. As the GEV is fitted to permuted data for any given data set, this failing means that the parameters estimated would differ from those of the hypothetical, 'true' null distribution. We found this to be the case when comparing GEV parameter estimates obtained from empirical and permuted data (Fig S13). Notable was the trend, or lack thereof, in the scale parameter: estimates from empirical data fell as the sample size increased, whilst those from permuted data showed no such trend. This behaviour was consistent with our observation of a smaller range of values in the empirical data at higher sample sizes, i.e. a reduction in the scale of the empirical distribution.

To confirm that the difference in estimated GEV parameters between empirical and permuted data could explain the downward trend in observed type 1 error rate for the GPS-GEV test, we tried estimating GEV parameters from empirical, not permuted, data. For each sample size, we fitted the GEV distribution to 100 realisations of the GPS statistic, computing each from a simulated replicate data set from the small-effect regime and with genetic correlation 0. As we had replicated each data set 200 times at each sample size, this left 100 replicate data sets at each sample size to form a 'test set' on which to evaluate the type 1 error rate obtained when using the GEV null distribution whose parameters were estimated from the first 100 data sets (Fig S14). Using only half of the replicate data sets in our test set yielded larger confidence intervals than those seen in the studies above, but the absence of a downward trend in type 1 error rate was clear.

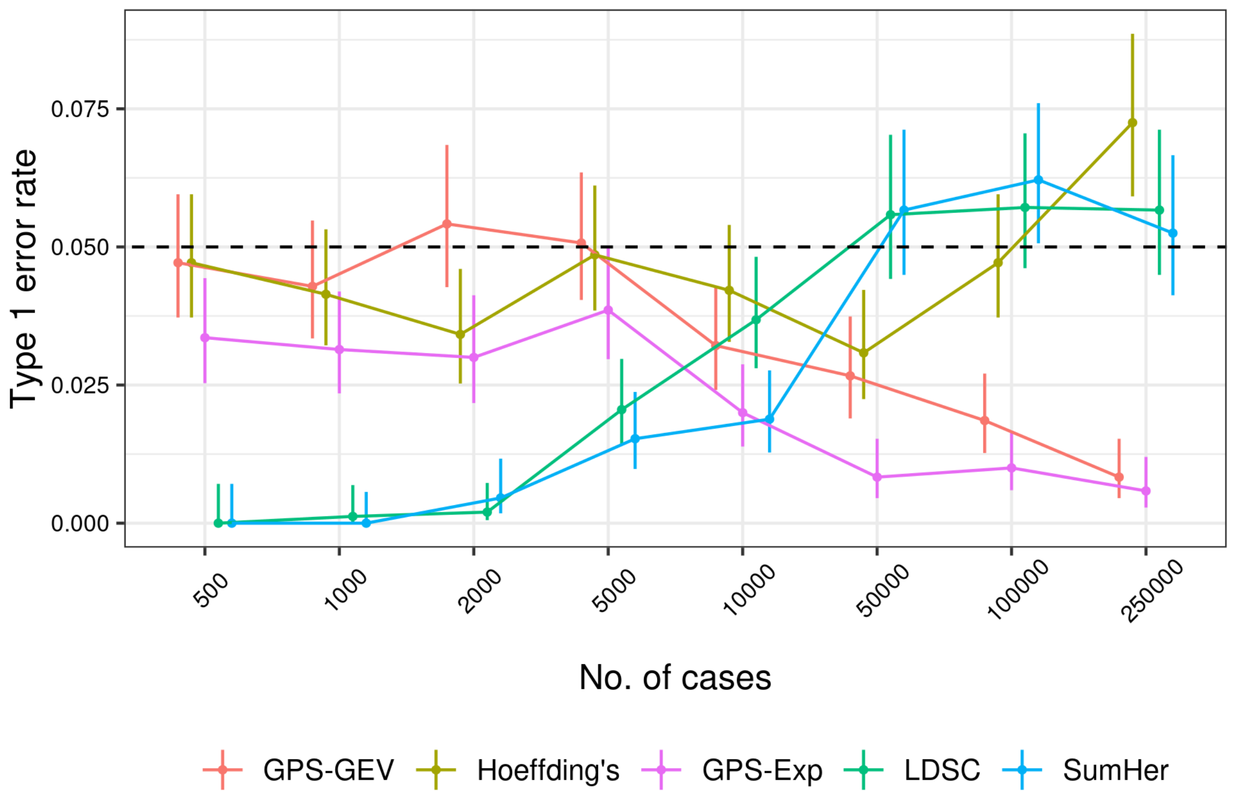


**Fig S11. The observed type 1 error rates of each method, inclusive of the additional 50,000 and 250,000 case and control samples.** The type 1 error rate for each method was estimated as the proportion of replicates for which $p \leq 0.05$ when the genetic correlation was set to zero. This proportion was measured only among those replicates of an initial 200 which produced a valid test statistic. 95% confidence intervals were calculated as Wilson score intervals. The dotted line depicts the size of the test, 0.05.

We next investigated the phenomenon of falling type 1 error rate for the GPS-Exp test, which makes no use of our permutation procedure. The GPS-Exp test instead uses the standard exponential as its null distribution: the rate parameter is held constant and is not estimated from the data, permuted or otherwise. As the GPS statistic is transformed to the reciprocal of its square to calculate its p-value for the GPS-Exp test (Equation 2), fewer large values on the untransformed scale would correspond to fewer small values on the transformed scale. Comparing empirical null data simulated with the smallest and largest sample sizes (500 and 250,000 cases, respectively), we found this to be the case: fewer (transformed) values fell below the critical value for a size $\alpha = 0.05$ GPS-Exp test at the highest sample size (Fig S15).

We were not able to devise a modification of our GPS-GEV permutation procedure to better recapitulate the behavior of empirical data, the consequence of which is a conservative test at very high sample sizes. We note this shortcoming is mitigated by the fact that a sample of 250,000 cases is an implausibly large number for a case-control study, even for a common disease study undertaken in the context of a modern biobank; the phenotype manifest of the Pan-UKBB GWAS collection shows the unitary disease phenotype with the largest number of cases to be ‘hypertension’ with 115,530, more than twice as many as the next largest, ‘high cholesterol’, with 54,681 (Text S2).

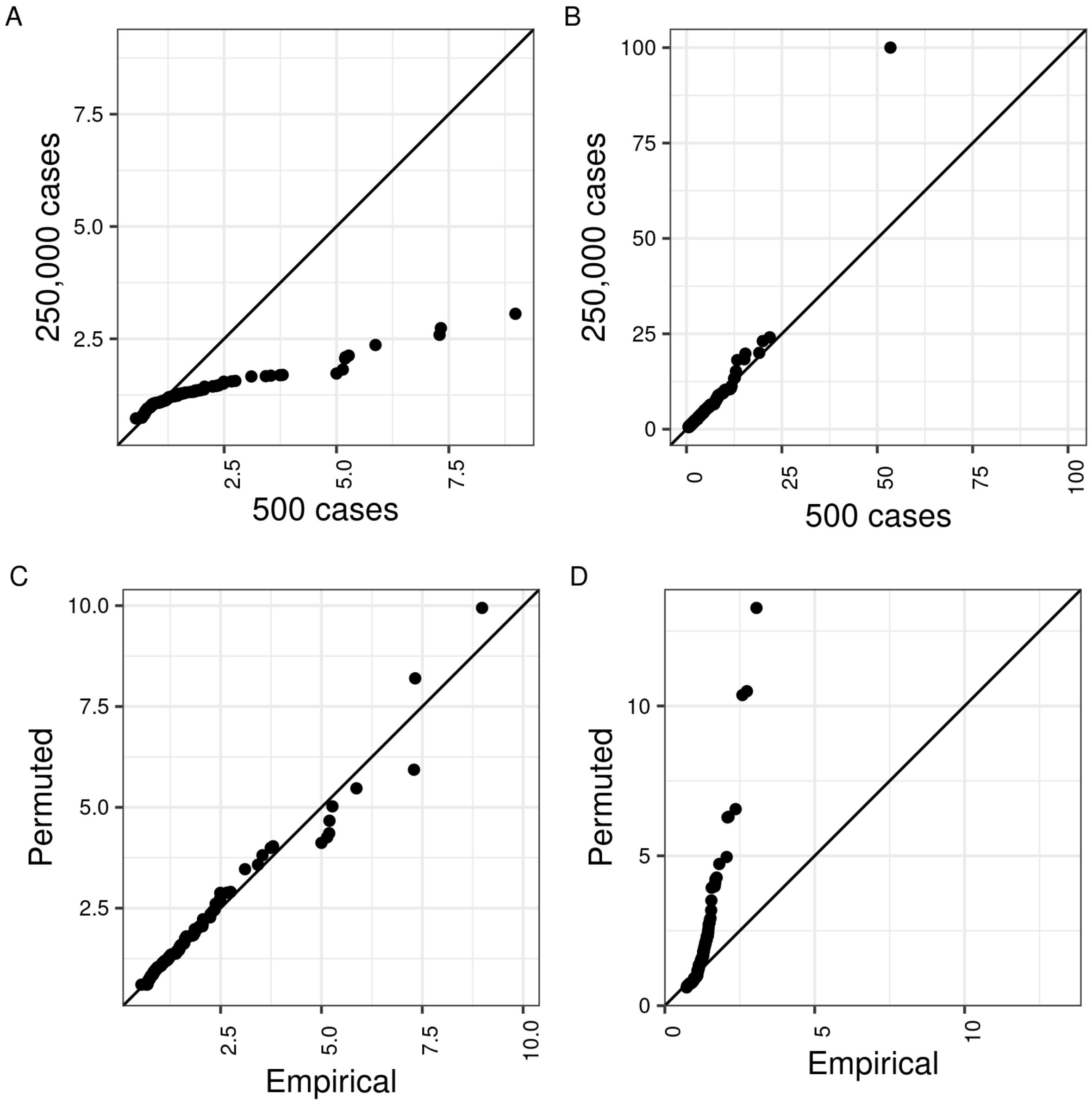


**Fig S12. The permutation procedure fails to reproduce the empirical null GPS distribution at higher sample sizes.** Two-sample quantile-quantile plots produced using data simulated under the small-effect regime. (A) GPS realisations from 200 replicate data sets simulated under the empirical null for the 500 and 250,000 case-control sample sizes; there are noticeably fewer large values in the 250,000 sample. (B) 200 GPS realisations generated through permutation of a single empirical null data set for the 500 and 250,000 case-control sample sizes. (C) 200 empirical and permuted null GPS realisations for the 500 case-control sample size. (D) 200 empirical and permuted null GPS realisations for the 250,000 case-control sample size; the empirical sample contains fewer large values than the permuted sample, which is not seen in (C).

We envision the use of the GPS-GEV test in the small-sample case, not the large, for reasons discussed in the main text. In the small-sample regime, we found that neither the GPS-Exp test

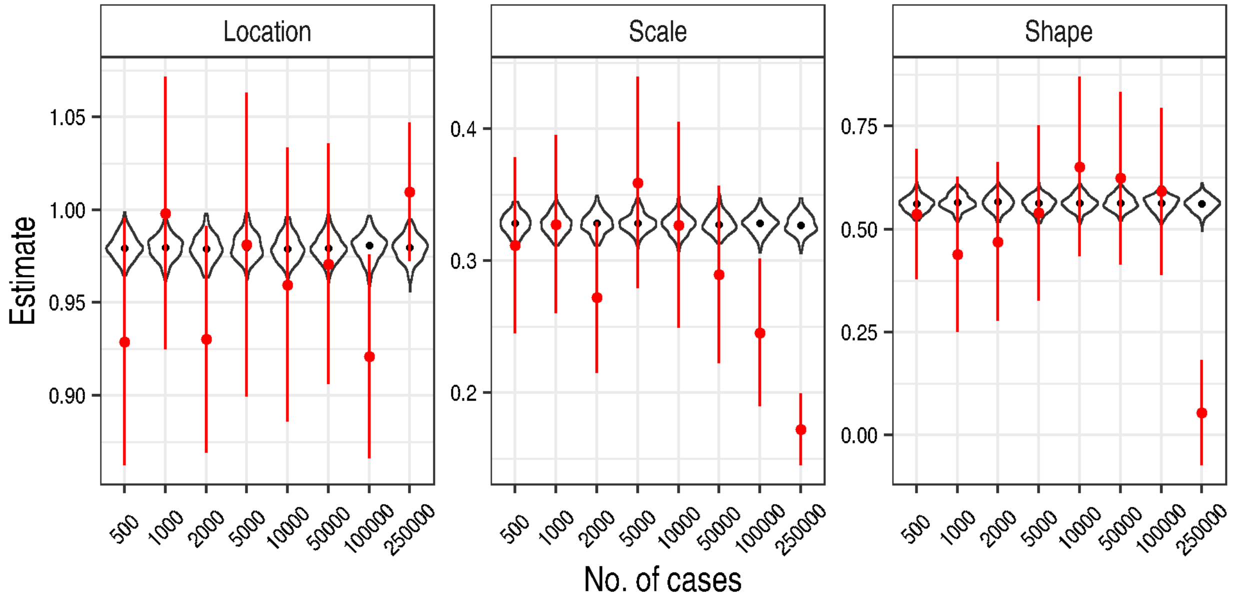


**Fig S13. Scale parameter estimates obtained from empirical, but not permuted, data fell with increasing sample size.** The distribution of parameter estimates as a function of sample size for the three parameters of the GEVD. Estimates were obtained from both empirical and permuted data from small-effect simulations with a genetic correlation of 0. The violin plots depict parameter estimates obtained from 200 replicate permuted data sets for each sample size; the black points depict the median of these estimates. The red points depict parameter estimates, one per sample size, obtained from 200 replicate empirical data sets. The red lines depict estimated 95% confidence intervals for the empirical estimates.

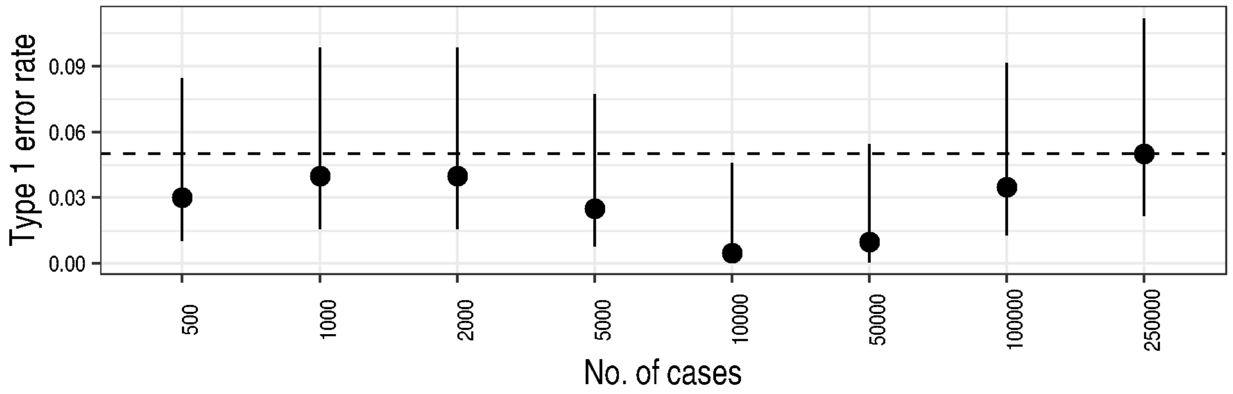


**Fig S14. Estimating GEV parameters from empirical, not permuted, data eliminates the downward trend in type 1 error for the GPS-GEV test**. The GPS statistic was computed for 200 replicate data sets simulated under the small-effect regime with a genetic correlation of 0 at each sample size. The first 100 realisations were used to fit the GEVD and the second 100 were used with the GPS-GEV test and the fitted distribution to measure the type 1 error rate. The type 1 error rate was estimated as the proportion of replicates for which $p \leq 0.05$. 95% confidence intervals were calculated as Wilson score intervals. The dotted line depicts the size of the test, 0.05.

nor the GPS-GEV test were overly conservative (Fig S11) and were in fact more powerful than their comparators for fewer than 5,000 cases and for five of six simulation regimes.

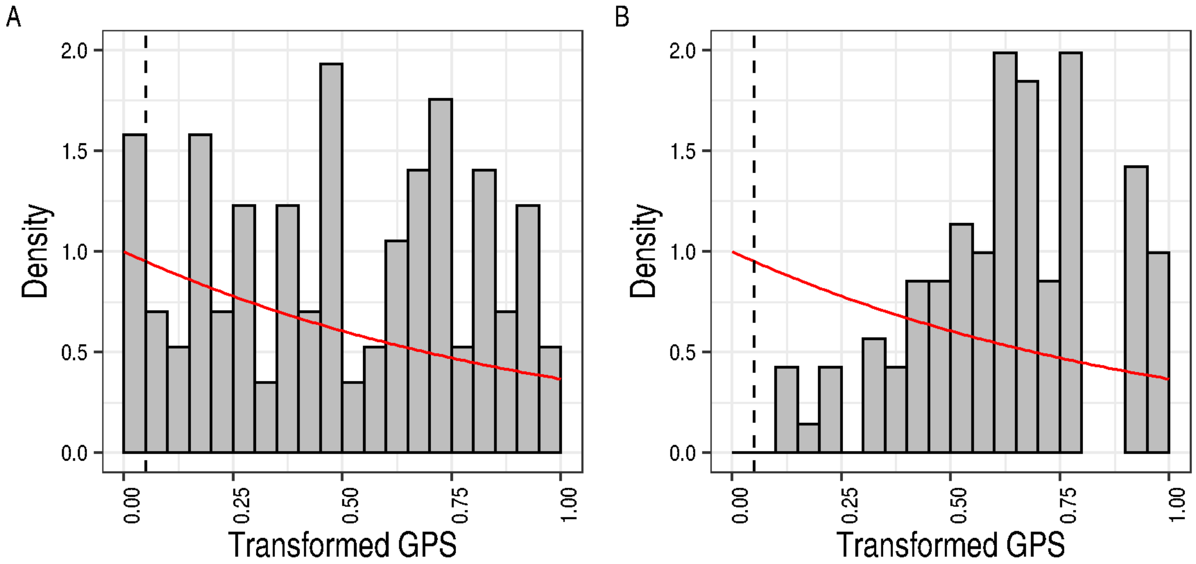


**Fig S15. A lack of very small test statistics on the transformed scale underlies the excessively conservative nature of the GPS-Exp test at the largest sample size.** Each subfigure depicts GPS test statistics computed from 200 simulated replicate data sets with a genetic correlation of 0 (i.e. under the empirical null) under the small-effect regime for (A) 500 cases and (B) 250,000 cases. The test statistics are on the transformed scale of the GPS-Exp test. The red line depicts the standard exponential density. The dashed vertical line depicts the critical value of the transformed statistic for the GPS-Exp test of size $\alpha = 0.05$; this value lies at the 5th percentile of the standard exponential distribution. The x-axis has been truncated to the $[0, 1]$ interval to more clearly depict the empirical distribution near the critical value.
